# COBREXA.jl: constraint-based reconstruction and exascale analysis

**DOI:** 10.1101/2021.06.04.447038

**Authors:** Miroslav Kratochvíl, Laurent Heirendt, St. Elmo Wilken, Taneli Pusa, Sylvain Arreckx, Alberto Noronha, Marvin van Aalst, Venkata P Satagopam, Oliver Ebenhöh, Reinhard Schneider, Christophe Trefois, Wei Gu

**Affiliations:** Luxembourg Centre for Systems Biomedicine, University of Luxembourg, Campus Belval, Belvaux, Luxembourg; Institute of Quantitative and Theoretical Biology, Heinrich-Heine-Universität Düsseldorf, Universitätsstraße 1, 40225 Düsseldorf, Germany; Nium S.à.r.l., 6 Avenue des hauts fourneaux, L-4362 Esch-sur-Alzette, Luxembourg; ELIXIR Luxembourg, 6, avenue du Swing, Campus Belval, L-4367 Belvaux, Luxembourg

## Abstract

COBREXA.jl is a Julia package for scalable, high-performance constraint-based reconstruction and analysis of very large-scale biological models. Its primary purpose is to facilitate the integration of modern high performance computing environments with the processing and analysis of large-scale metabolic models of challenging complexity. We report the architecture of the package, and demonstrate how the design promotes analysis scalability on several use-cases with multi-organism community models.

**Availability and implementation:** https://doi.org/10.17881/ZKCR-BT30.

## 1 Introduction

Understanding metabolic interactions in cells is a crucial step to investigate disease mechanisms and to discover new therapeutics (Cook and Nielsen, 2017; Apaolaza et al., 2018; Brunk et al., 2018). Constraint-Based Reconstruction and Analysis (COBRA) is a promising methodology for analyzing various metabolic processes at the organism- and community-levels (Fang, Lloyd, and Palsson, 2020). The main idea behind COBRA is to represent an organism as a constrained set of inter-connected reactions and metabolites based on genomic sequencing data. This leads to a straightforward interpretation of metabolism as a constrained linear system, which enables the utilization of a wide range of well-developed analysis methods (Orth, Thiele, and Palsson, 2010).

The increasing ubiquity of genomic sequencing has led to a rapid expansion in the number and complexity of genome-scale metabolic models, e.g. the human metabolic model that has more than 80,000 reactions (Thiele et al., 2020). Recent automated reconstruction tools can generate models spanning the entire primary metabolism of both pro- and eukaryotes (Machado et al., 2018). Consequently, metabolic models are becoming considerably larger in scale than their predecessors, which is further compounded by the construction of multi-member community models. This growth implies increasing analysis complexity (see Figure S1), which in turn drives the need to develop analysis software that can accommodate this complexity. While computing the solutions to the underlying constrained optimization problems is hard to accelerate and parallelize, many analysis types can be decomposed into individual invocations of the optimizer, which may be parallelized. However, despite continued efforts (Heirendt, Thiele, and Fleming, 2017), this remains challenging due to the scalability limits of existing software implementations.

Here, we present COBREXA.jl, a package for implementing and running distributed COBRA workflows. The package is implemented in the Julia programming language (Bezanson et al., 2017), enabling facile extension with user-defined numeric-computing routines, and interoperability with many high-performance computing packages. It provides a ‘batteries-included’ solution for scaling analyses to make efficient use of high-performance computing (HPC) facilities, giving researchers a powerful toolkit for executing complicated high-volume workflows, such as the creation and exploration of digital metabolic twins in personalized medicine (Björnsson et al., 2020), and analysis of extensive microbial communities in ecology and biotechnology. We report the implementation architecture, and substantiate how the design accommodates future extensions and scaling of common analysis tasks.

## 2 Implementation and results

COBREXA.jl is an open architecture solution, providing interchangeable building blocks for implementing complicated COBRA workflows. Common analysis methods, such as flux balance, flux variability, and gene knockout analyses (Gudmundsson and Thiele, 2010), are implemented as ready-to-use functions that may be easily composed and customized. Most importantly, the building blocks are designed so that the constructed workflows can be easily separated into parallelizable analysis steps and executed on multiple computation nodes in HPC environments (as illustrated in Fig. 1). The concurrent execution of such workflows results in significant computational speedups, without requiring user expertise in parallel programming.

**Figure 1:**
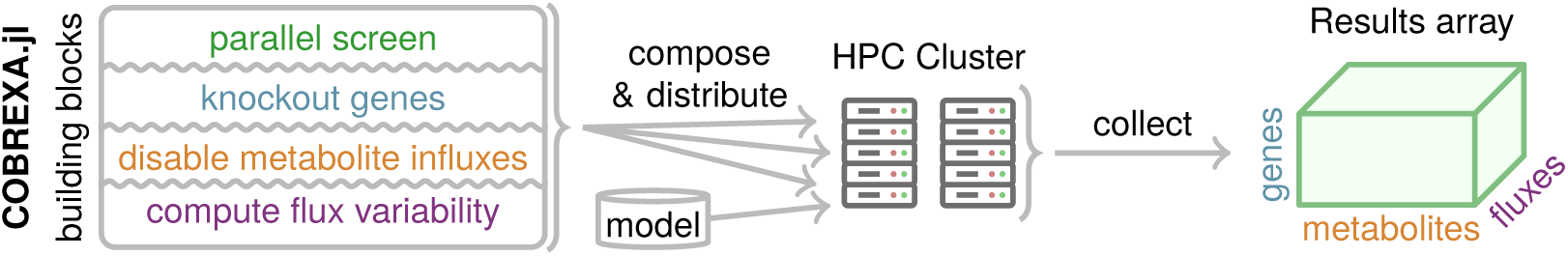
Schema of an example custom analysis construction that examines flux variability in many variants of a model, its distributed execution with COBREXA.jl, and collection of many results in a multi-dimensional array.

The design of COBREXA.jl distinguishes it from other COBRA implementations, which typically provide parallelization support for only a few selected methods, and no current support for parallelization of custom method variants. For example, parallel single-gene deletion analysis is commonly supported, but a variant that explores the flux variability in knockouts must be reimplemented and parallelized by the user.

A variety of model exchange and representation formats are supported, including MATLAB format (Heirendt, Arreckx, et al., 2019); object-oriented JSON format (Ebrahim et al., 2013), and SBML (Keating et al., 2020). Additionally, implementation of the workflows in Julia results in highly optimized execution of the code at the cost of minor pre-compilation overhead, which benefits large, data-heavy use cases. A detailed architecture overview is provided in Supplementary Section S1.

To evaluate the effect of the new architecture and optimizations on the performance and scalability of COBRA analyses, we benchmarked COBREXA.jl on use-cases that benefit from parallelization. We compared its performance to that obtained with COBRApy (Ebrahim et al., 2013) and COBRA Toolbox (Heirendt, Arreckx, et al., 2019), which are the widely adopted tools for running COBRA workflows. Running on a 256-CPU multi-node cluster, COBREXA.jl was able to fully utilize the available distributed computing resources and outperform the implementation of flux variability analysis in other packages by a factor of between 2× and 10×, even on relatively small models (Supplementary Table S2). We further demonstrated that COBREXA.jl is able to parallelize and distribute custom workloads by re-implementing the production envelope functionality of COBRApy; leading to speedups of over 10×, even on a single 16-core computation node (Supplementary Table S3). Consequently, we expect that the COBRA methods implemented in COBREXA.jl will enable reliable acceleration of many current and future workloads by simply adding more computing resources. The results are further discussed in Supplementary Section S3.4.

## 3 Conclusion

COBREXA.jl is a new package developed for large-scale distributed processing of constraint-based biological models. It differs from the other implementations of COBRA methods (Heirendt, Arreckx, et al., 2019; Ebrahim et al., 2013) by focusing on computational efficiency, and simplifies high-level construction of parallelized user-defined analysis methods. This is required for performing extensive analyses of large models, future-proof extensibility, and workload distribution that enables effective utilization of the common HPC infrastructure resources. The package thus enables fast analysis of datasets that may pose challenges for the currently available tools, such as the comprehensive human gut microbiome models.

