## Supplementary information for "COBREXA.jl: constraint-based reconstruction and exascale analysis"

### S1 Internal structure of COBREXA.jl

This section summarizes the design rationale and extensibility of the COBREXA.jl package.

#### S1.1 Polymorphic models future proofs analysis workflows

COBREXA.jl does not enforce a single universal model type. This ensures that the package will be able to incorporate novel model structures that are possibly incompatible with current standards and assumptions, while maintaining the functionality of the existing analysis workflows. To ensure that the new extensions incorporate well into the package ecosystem, COBREXA.jl defines a structure-agnostic interface for accessing the model data and enforces its use in the main analysis functions. In the long term, this promotes separation of concerns and avoids the feature creep that inevitably occurs in any all-encompassing data structure.

As a typical example, new structure of the annotations and constraints may be required to be implemented in the models in the future, possibly causing incompatibility with any current data layout; separating the new model structure prevents the changes from breaking the existing code. Custom model data structures may also be designed to utilize specific optimizations for particular use-cases without the rist of causing undesirable overhead in others – for example, implementing a model with index structures that allow rapidly computing knockouts of reactions by selected genes will not impose the re-indexing overhead in models and analyses that do not require work with genes.

The custom model structures interface with COBREXA.jl analysis functions using a polymorphic interface of several overloaded accessor methods, which rely on multiple dispatch system of Julia. The accessors methods include simple functions to recreate the stoichiometric matrix, bounds and objective vectors, and functions that access various model annotations (if available), including the gene-reaction relationships and chemical formulas of the metabolites. The analysis functions use only these generic accessors to read the necessary information from models, which ensures their portability to any future model structure implemented by the users.

COBREXA.jl includes ready-to-use model types that reflect the common practice in current metabolic modeling, including the type `CoreModel` (a COBRA Toolbox-style sparse matrix model), `StandardModel` (an object-oriented-style model similar to COBRApy models), and `SBMLModel` (which mimics the structure of Systems Biology Markup Language as described by Keating et al., 2020). Thanks to the polymorphic design, all model types are freely interchangeable in the implemented workflows, and can be supplemented by the user by implementing another subtype of the abstract `MetabolicModel` type.

An up-to-date description of the model interface can be obtained from the documentation at `https://lcsb-biocore.github.io/COBREXA.jl/`. An example implementation of the accessor structure for a very simple custom model used in benchmarking can be seen in supplementary script `benchmark-prep.jl` (Table S1).

#### S1.2 Specification of model variants and solver modifications

To allow for straightforward implementation of large analyses, COBREXA.jl utilizes the high-level programming capabilities of Julia to create a set of model and analysis modifiers that can be used to declaratively specify various properties of the analysis sub-tasks. In the distributed processing framework, these functions act as small 'recipes' that are efficiently transferred to remote workers where they make larger changes to the model structure. All analysis functions accept a list of such modifiers and apply them to the model before the analysis, providing a useful way for e.g. controlling specific optimizer parameters without any additional implementation overhead.

For example, flux balance analysis may be modified declaratively to solve a different objective (declared by `set_objective`) with one reaction knocked out (declared by `set_bounds`) as follows:

```
optimal_flux = flux_balance_analysis_vec(
    model,  # a loaded model
    Tulip.Optimizer;  # the linear problem solver to use
    modifications = [
        set_objective("Reaction1"),  # tell the optimizer to optimize the rate of Reaction1
        set_bounds("Reaction2", 0, 0),  # set the lower and upper bound of Reaction2 to 0
    ]
)
```

Specific functions are provided to run parallel analysis of many model variants at once. That way, with a model type that supports the gene knockouts, one can easily combine the basic tools to run a flux balance analysis for all combinations of single-gene knockouts and single-reaction knockouts, producing a matrix of the objective values from all sub-results as follows:

```
results = screen(
    model;
    variants = [ # a matrix of all variants to be examined
        [
            # each variant is specified by a list of the modifications:
            with_gene_knockout(g),
            with_removed_reaction(rxn)
        ]
        for rxn=reactions(model), g=genes(model)
    ],
    # the `analysis` is a function executed independently on each variant
    analysis = m -> objective_value(flux_balance_analysis(m, Tulip.Optimizer)),
    # parallelism is enabled by specifying IDs of the worker processes:
    workers = [2,3,4,5]
)
```

##### S1.3 Model distribution optimizations

COBREXA.jl uses the DistributedData.jl package (`https://github.com/LCSB-BioCore/DistributedData.jl`, Kratochvíl et al., 2020) for explicitly caching temporary copies of frequently reused dataset objects (in most cases, the whole model structure) on remote workers, avoiding the potentially slow transfer of base data during the distributed analysis. Distributed and parallel processing is enabled by passing a worker list to any given analysis function.

The model distribution process is optimized to utilize shared storage, which is a common (although not standardized) feature of many HPC systems. To allow the users to benefit from this feature while avoiding portability problems (such as HPC-system-specific implementations of analysis functions), the feature is enabled by utilizing a simple 'wrapper' model structure that transparently serializes any other model to a fast shared storage and loads it on demand on remote workers. The specific disk-cached type, called `Serialized{M}`, works with any subtype `M` of `MetabolicModel`. The benchmark script `benchmark-prep.jl` (Table S1) demonstrates the use with a simple custom model type.

#### S2 The need for scalable COBRA implementations

To illustrate the scale of computation required for various classes of realistic, biologically relevant problems, we catalogued the size of existing published unicellular models (collected from AGORA and BIGG databases), several multicellular organism models, whole-organ and whole-body reconstructions, and various publications referencing the microbiomes of human, bovine, canine, ocean, and soil. Figure S1 shows

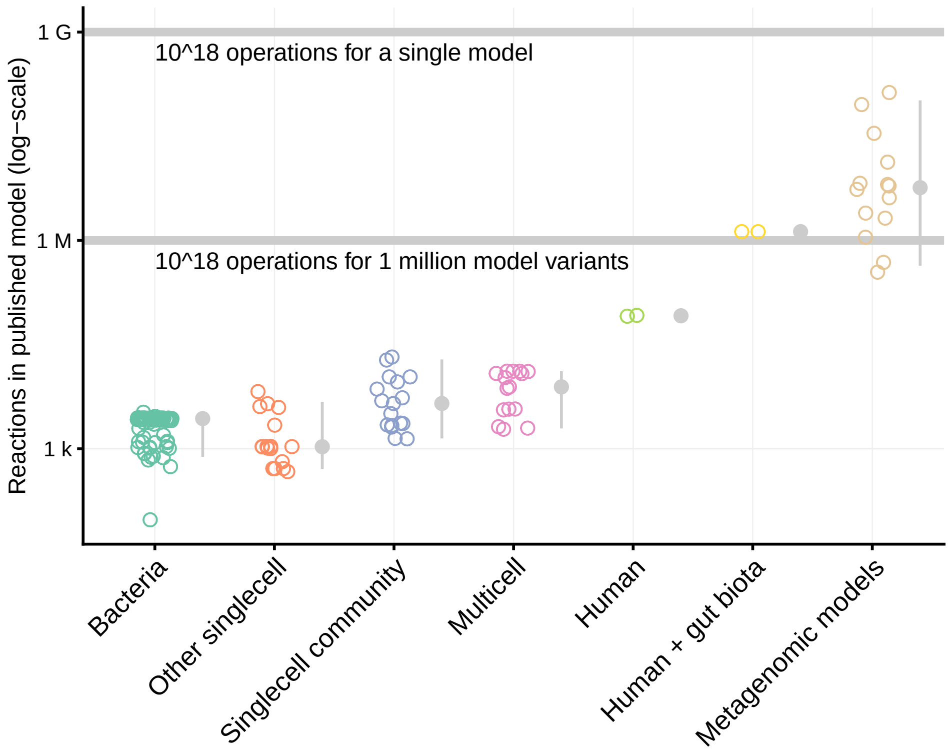


Figure S1: Size of the published models (represented by dots) can easily cause the analysis to reach huge computation volumes. The latest human microbiome models would cross the threshold of $10^{18}$ required operations (one 'exa-operation') if around 4 million model variants would be processed using typical algorithms, such as FVA. Models constructed from metagenomic data exceed this bound at mere thousands of analyzed variants. The model size thresholds are highlighted by horizontal lines.

the number of reactions contained in each system (putative in the case of the metagenomic systems). Typically, analysis of metabolic models combines a constraint-based analysis with generation and scrutinization of multiple variants of the model or the optimization objective (e.g., flux balance analysis on multiple gene and organism knockouts, flux variability analysis on all reactions, ...). In Figure S1, the markers illustrate the putative model size where the usual analysis will inevitably reach $10^{18}$ required basic operations i.e. the exa-scale boundary. For the estimation, we considered a flux-variability-style analysis on all model reactions, which requires at least $\mathcal{O}(|\text{reactions}|^2)$ computation steps in the case of a single model, possibly multiplied by the total number of examined model variants. Notably, the notion of "operations" used here is asymptotic – actual instruction count on realistic hardware will be much higher, thus further lowering the thresholds.

### S3 Performance and scalability on biological use cases

To assess the scalability and applicability of COBREXA.jl, we performed two benchmarks on realistic use-cases that demonstrate the scalability results and can be used to estimate the viability of using the different COBRA implementations for computational tasks of various sizes. An additional benchmark was performed to estimate the impact of data distribution optimization in the HPC environments.

#### S3.1 Benchmarked datasets

We used the human gut microbiota models from AGORA database. Pseudorandomly generated subsets of all microbes were combined into small community models as follows: Exchange reactions in the individual models were replaced by reactions that transfer the metabolites between the extracellular compartment of the model and the "community supply" of the metabolite, represented as a special metabolite. Biomass exchange reactions were processed in the same way, connected with a forward-only reaction that adds the biomass into the community biomass pool. Additional reactions were added that provide exchange of the community metabolite supply with the environment (i.e., supply or remove each metabolite from the system). These were constrained to allow only a small influx of the metabolites compared to outflux (lower bound $= -1000$, upper bound $= 10$), except for water, oxygen and carbon dioxide where both large influx and outflux was allowed (upper bound $= 1000$). Additional unidirectional reaction for removing the community biomass from the model was added, and set as the objective of the model. Using this procedure, we generated models that contained 5, 10, 20, and 50 individual microbes, each with several differently seeded subsets of the AGORA microbes. Given the average size of a single microbe model of around 1800 reactions, this allowed us to easily generate biologically meaningful models of many required sizes. The details are available in supplementary script `benchmark-prep.jl` (Table S1).

#### S3.2 Benchmarked use-cases

We designed two benchmarks that substantiate the benefits of using COBREXA.jl stated in main text Section 2. The first benchmark measures the speedup that COBREXA.jl can gain by utilizing distributed workers, as compared to a package that does not implement workload distribution. The second benchmark shows that the parallelization speedup is available also for newly implemented and user-specified analyses that are typically not parallelized in the existing COBRA software.

**Benchmark 1: Flux variability analysis** We chose flux variability analysis as the main benchmark for determining the scalability of the packages, due to wide availability of parallelized implementations. The corresponding functions from the packages (`flux_variability_analysis` in both COBRApy and COBREXA.jl) were called to resolve the variability of all reactions in the model with the objective value bound set to 90% of the global objective. The details can be found in scripts `benchmark-fva.jl` and `.py` (Table S1).

**Benchmark 2: Production envelopes** We chose the computation of production envelopes in the model as an example of an analysis that can be easily parallelized and can be constructed from the building blocks in COBREXA.jl, but many packages do not implement the parallelization, forcing the users to fall back to the serial implementations. The production envelopes were created for 3-dimensional grids in the flux space. We selected the dimensions from the available exchange reactions that supply energetically interesting metabolites from the environment (choice of the exchanges was made from first 3 exchanges of sucrose, L-glutamine, galactose, L-glutamate, D-glucose, xylose and arabinose, in order, that were present in the model). The flux bounds of the selected 3 reactions were set to small influx of the metabolite (bounded between 0 and 100). The envelope computation was then benchmarked for 7, 10, and 14 points in each dimension (respectively translating into 343, 1000 and 2744 individual optimization problem instances to solve). Only the reachable flux values were collected. Details are available in scripts `benchmark-envelopes.jl` and `.py` (Table S1).

#### S3.3 Benchmark setup

**Computation environment** The benchmarks were conducted on the hardware of the 'iris' cluster of University of Luxembourg HPC facility, see `https://hpc.uni.lu/`. The computational nodes were equipped with Intel Xeon E5-2680 v4 @2.4GHz CPUs with 128GB RAM per node; nodes were interconnected with 100GB/s Infiniband EDR. All benchmarks were executed using the Slurm scheduler, using the scripts in Table S1. Measurements were collected from the log files generated by the scripts.

Measurements of COBRA Toolbox performance were collected on different hardware due to software compatibility and licensing issues; we used a single computation node with Intel Intel Core i5-1038NG7 @2.00GHz and 16GB of RAM. The absolute scales of the results may differ by a constant factor; single-core performance of i5-1038NG7 is estimated 21% higher (precise estimate of the CPU performance differences can be seen e.g. at `https://www.cpubenchmark.net/compare/Intel-i5-1038NG7-vs-Intel-Xeon-E5-2680-v4/3723vs2779`). This does not affect the interpretation of our benchmark results, which aim to show mainly the impact of multi-core parallelization.

**Software** We used COBREXA.jl version 1.0.5 (git commit `d55323dd` from `https://github.com/LCSB-BioCore/COBREXA.jl`, with Julia version 1.6.2 and JuMP.jl interface version 0.21.9), COBRApy version 0.22.1 (Ebrahim et al. (2013), installed from `pip`, running on Python 3.8.6 with optlang interface version 1.5.2), and COBRA Toolbox version 3.33 (running on MATLAB 2020b). For solving the constrained linear problems, we always utilized the Gurobi optimizer version 9.1. In all cases, we only enabled the parallelization methods that were directly supported and documented by the packages, i.e., we did not attempt to improve the parallelization with external scriptage or manual distribution of the analysis execution to worker nodes.

The reported measurements are averaged from multiple runs. In case of COBREXA.jl a 'dummy' run with minimal data was executed before collecting benchmark results to allow precompilation of Julia code. The benchmarks were repeated for multiple model variants of similar size but different contents (differently seeded random choices of the models used to construct the community, see Section S3.1) to compensate for the effect of data-dependent solver performance.

#### S3.4 Benchmark results and discussion

Results from benchmark 1 in Table S2 show that COBREXA.jl is able to continue to scale to large analysis and problem sizes and benefit from any available distributed computing resources. In particular, we observed that COBREXA.jl performs similarly to the state-of-the-art packages in situations where the resources fit on a single computation node, but continues to scale even when the available resources cannot fit on a single node, and thus cannot be utilized by shared-memory parallelism schemes. That demonstrates a common limit in the COBRA packages that can otherwise be only partially countered by the users, usually by implementing custom parallelization and external task distribution frameworks.

In the benchmark we also observed parallelization overhead in COBREXA.jl, especially with the smaller models and higher node counts, but the overhead became negligible for larger analyses. Notably, the scaling advantage was substantial even for the relatively small models of 50 organisms. The observations about scalability are also supported by the previous benchmarking the code of package `DistributedData.jl`, showing that it is able to scale to much larger and more communication-intensive use-cases (see Supplementary Figure S2 in the report of Kratochvíl et al. (2020) for details). Based on the benchmarks, we expect COBREXA.jl to be able to reliably utilize any added computing resources to provide substantial parallelization speedups for growing model sizes.

As a side benefit of using a compiled language, we observed a surprisingly high speedup of COBREXA.jl operations that do not directly depend on the performance of the solver but rather on the performance of the programming environment, such as the model loading (Figure S2).

This is further confirmed by results from benchmark 2 (Table S3) even for the analysis methods constructed from the building blocks provided by COBREXA.jl, which possess no parallelized implementation in other packages. Notably, COBREXA.jl implementation of the production envelopes is constructed from a generic function for parallel screening through a parameter space, which is incidentally the same as the one used to implement the flux variability analysis.

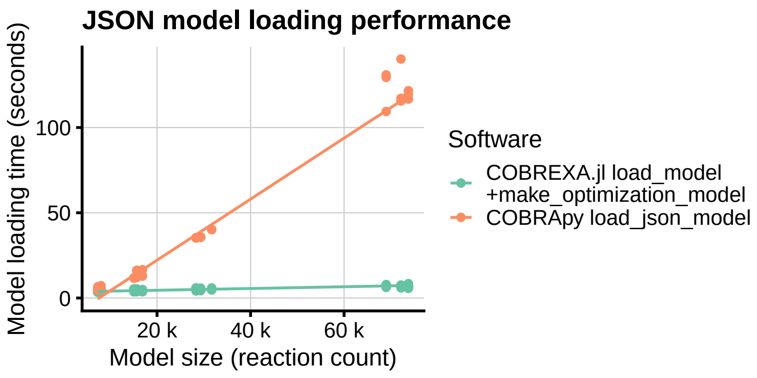


Figure S2: Time required to load a JSON model by COBRA implementations. For efficiency reasons, COBREXA.jl model loading functions do not create an optimizer instance of the model automatically upon load as does COBRApy. This was compensated in the benchmark by explicitly building the optimizer instance in COBREXA.jl after loading, using function `make_optimization_model`.

| File | Contents |
|---|---|
| `gen.sh` | Shell generator of Slurm batch scripts that can be scheduled using the `sbatch` command. Scheduling the batch scripts then indirectly executes the Julia and Python scripts. |
| `benchmark-prep.jl` | Julia script for preparing the base community models for the benchmarks. |
| `benchmark-fva.jl` | FVA benchmark (COBREXA.jl version). |
| `benchmark-fva.py` | FVA benchmark (COBRApy version). |
| `benchmark-fva.m` | FVA benchmark (COBRA Toolbox version). |
| `benchmark-envelopes.jl` | Envelope computation benchmark (COBREXA.jl version). |
| `benchmark-envelopes.py` | Envelope computation benchmark (COBRApy version). |

Table S1: Overview of the supplementary benchmark scripts, available at `https://doi.org/10.17881/ZKCR-BT30`.

| | Model size | 1 core | 4 cores | 16 cores | 64 cores | 256 cores | Speedup |
|---|---|---|---|---|---|---|---|
| COBRApy | 5 | 125.39 | 53.05 | 18.02 | – | – | 1× |
| | 10 | 595.54 | 257.92 | 93.43 | – | – | |
| | 20 | 2539.34 | 1310.42 | 415.5 | – | – | |
| | 50 | × | 9421.9 | 2887.81 | – | – | |
| COBREXA.jl | 5 | 131.57 | 35.6 | 12.04 | **9.56** | × | 1.88× |
| | 10 | 939.28 | 282.1 | 76.82 | **35.8** | 64.96 | 2.61× |
| | 20 | 4862.36 | 1465.15 | 477.05 | 164.24 | **139.87** | 2.97× |
| | 50 | × | 10818.09 | 3618.13 | 1066.48 | **584.37** | 4.94× |
| COBRA Toolbox* | 5 | 368.97 | 143.14 | – | – | – | – |
| | 10 | 1787.95 | 942.94 | – | – | – | |
| | 20 | 14520.76 | 7093.15 | – | – | – | |
| | 50 | × | × | – | – | – | |

Table S2: Results collected from benchmark 1 show dependency of the computation time (in seconds) of the FVA analysis on the available CPU core count (see Section S3.4 for discussion). Speedups are computed between the best results for a given model size. Cases where the required multi-node distribution or parallelization was not available are marked with '–'. Corner cases (marked with '×') were not executed for resource efficiency. Measurements of COBRA Toolbox (marked with *) were collected on different hardware (see Section S3.3).

| Model size | Package | CPU cores | Envelope computation time (seconds) $7^3$ samples | $10^3$ samples | $14^3$ samples |
|---|---|---|---|---|---|
| 5 | COBRApy | 1 | 8.69 | 16.8 | 48.91 |
| | COBREXA.jl | 1 | 6.67 | 12.59 | 26.84 |
| | | 4 | **5.11** | 8.41 | 13.11 |
| | | 16 | 6.76 | **6.91** | **10.4** |
| 10 | COBRApy | 1 | 23.97 | 51.24 | 99.88 |
| | COBREXA.jl | 1 | 17.67 | 32.25 | 71.64 |
| | | 4 | **12.25** | 20.49 | 32.33 |
| | | 16 | 13.35 | **16.27** | **20.19** |
| 20 | COBRApy | 1 | 78.49 | 153.73 | 269.56 |
| | COBREXA.jl | 1 | 43.92 | 77.1 | 161.48 |
| | | 4 | 31.08 | 61.88 | 122.77 |
| | | 16 | **31.03** | **40.93** | **58.77** |
| 50 | COBRApy | 1 | 1100.35 | 1386.66 | 1768.12 |
| | COBREXA.jl | 1 | 241.75 | 433.49 | 788.14 |
| | | 4 | 155.35 | 233.87 | 470.66 |
| | | 16 | **106.86** | **155.38** | **222.48** |

Table S3: Speed of solving the model variants required for a 'production envelope' analysis. The trivial implementation built from the components available in COBREXA.jl is able to scale with added computing resources, making the method feasible for analysis of large models.
